# Cdk activity drives senescence from G2 phase

**DOI:** 10.1101/041723

**Authors:** Erik Müllers, Helena Silva Cascales, Libor Macurek, Arne Lindqvist

**Author notes:** Corresponding author: Arne Lindqvist Department of Cell and Molecular Biology Karolinska Institutet Stockholm, Sweden.

## Abstract

In response to DNA damage a cell can be forced to permanently exit the cell cycle and become senescent. Senescence provides an early barrier against tumor development by preventing proliferation of cells with damaged DNA. By studying single cells, we show that Cdk activity is retained after DNA damage until terminal cell cycle exit. The low level of Cdk activity not only allows cell cycle progression, but also forces cell cycle exit at a decision point in G2 phase. We find that Cdk activity stimulates p21 production, leading to nuclear sequestration of Cyclin B1, subsequent APC/C^Cdh1^-dependent degradation of mitotic inducers and induction of senescence. We suggest that the same activity that triggers mitosis in an unperturbed cell cycle drives senescence in the presence of DNA damage, ensuring a robust response when most needed.

## Introduction

In response to DNA damage, the cell cycle is halted to allow DNA repair. This is particularly critical in G2 phase, as entry into mitosis with unrepaired DNA may result in chromosomal aberration and propagation of mutations. Several mechanisms that establish a G2/M arrest by counteracting mitosis-promoting factors have been described [1–3]. However, upon DNA damage in S or G2 phase the production of mitosis-inducing factors such as Cyclin A2, Cyclin B1, Aurora A, Aurora B and Plk1 initially continues, albeit at a reduced pace [4–6]. As multiple feedback loops ensure a spiraling activation of Cyclin B1-Cdk1, Cyclin A2-Cdk1/2 and Plk1 that ultimately results in mitotic entry [7], maintaining even low levels of mitosis-inducing factors poses the risk to eventually overrun a cell cycle arrest. In fact, suppression of Cdk activity merely by posttranslational modifications is insufficient to sustain a G2/M arrest [8–10].

To avoid override of a cell cycle arrest, cells have evolved mechanisms that force terminal cell cycle exit and senescence. We and others have shown that terminal cell cycle exit from G2 phase depends on p53, its transcriptional target p21, and activation of the ubiquitin ligase APC/C^Cdh1^ that targets a large number of cell cycle regulators for degradation [5,6,11–13]. During this process, cells lose the expression of G2-specific proteins, exit the cell cycle, and become senescent, thereby preventing propagation of mutations [11,13–16]. What determines whether and when a cell in G2 phase becomes senescent remains unclear.

There are several indications that induction of senescence is a regulated process. Terminal cell cycle exit is a sharp transition, whose point-of-no-return is marked by the translocation of Cyclin B1 from the cytoplasm to the nucleus [6,11,17]. Before terminal cell cycle exit is initiated in G2 phase there is a variable delay, whose duration depends on when within S or G2 phase the damage occurred. That is, a cell receiving damage in late G2 exits the cell cycle faster than a cell receiving damage in early G2 phase [6]. These observations suggest that the signaling pathways that mediate senescence and cell cycle progression are interlinked.

Here we show that, despite a severe suppression, Cdk activity is sustained during a cell cycle arrest until terminal cell cycle exit occurs and that this remaining Cdk activity is needed to promote timely induction of senescence. We suggest that the key activity that drives mitosis in the absence of DNA damage drives cell cycle exit in the presence of DNA damage.

## Results

### Concerted Cdk1 and Cdk2 activity regulates Cyclin B1 nuclear translocation and induction of senescence upon DNA damage

We previously employed live-cell microscopy of individual RPE cells encoding a Cyclin B1-eYFP fusion protein at the endogenous *CCNB1* locus to study terminal cell cycle exit. Using this system we observed that DNA damage-dependent nuclear translocation and degradation of Cyclin B1 occurred only after S-phase completion [6]. As this is the time when Plk1 and Cdk1 are activated in an unperturbed cell cycle [18], we hypothesized that cell cycle kinases might be involved in regulating cell cycle exit after DNA damage. To test this idea we monitored the effect of different cell cycle kinase inhibitors on Cyclin B1-eYFP levels and localization after DNA damage using time-lapse microscopy.

To study cell cycle-dependent effects without affecting initiation of the DNA damage response (DDR), we added small molecule inhibitors of cell cycle kinases 1h after addition of the topoisomerase inhibitor Etoposide. Addition of a potent inhibitor of Plk1 did not affect Cyclin B1-eYFP degradation or nuclear translocation upon DNA damage, indicating that Plk1 is not required for these events (Supplementary Fig 1A). On the contrary, addition of selective Cdk1 or Cdk2 inhibitors alone led to a limited delay in the onset of Cyclin B1-eYFP nuclear translocation. Moreover, combined addition of selective Cdk1 and Cdk2 inhibitors, or addition of the broad Cdk inhibitor Roscovitine [19] almost completely abolished nuclear translocation of Cyclin B1-eYFP and delayed the onset and duration of Cyclin B1-eYFP degradation (Fig 1A-C and Supplementary Fig 1B and D). Similarly, stimulation of cellular Cdk activity by inhibition of Wee1 led to earlier degradation along with higher rates of DNA damage checkpoint slippage (Supplementary Fig 1C). Taken together, our results suggest that Cdk1/2 activity is needed for timely Cyclin B1 translocation and degradation after DNA damage.

**Figure 1.**
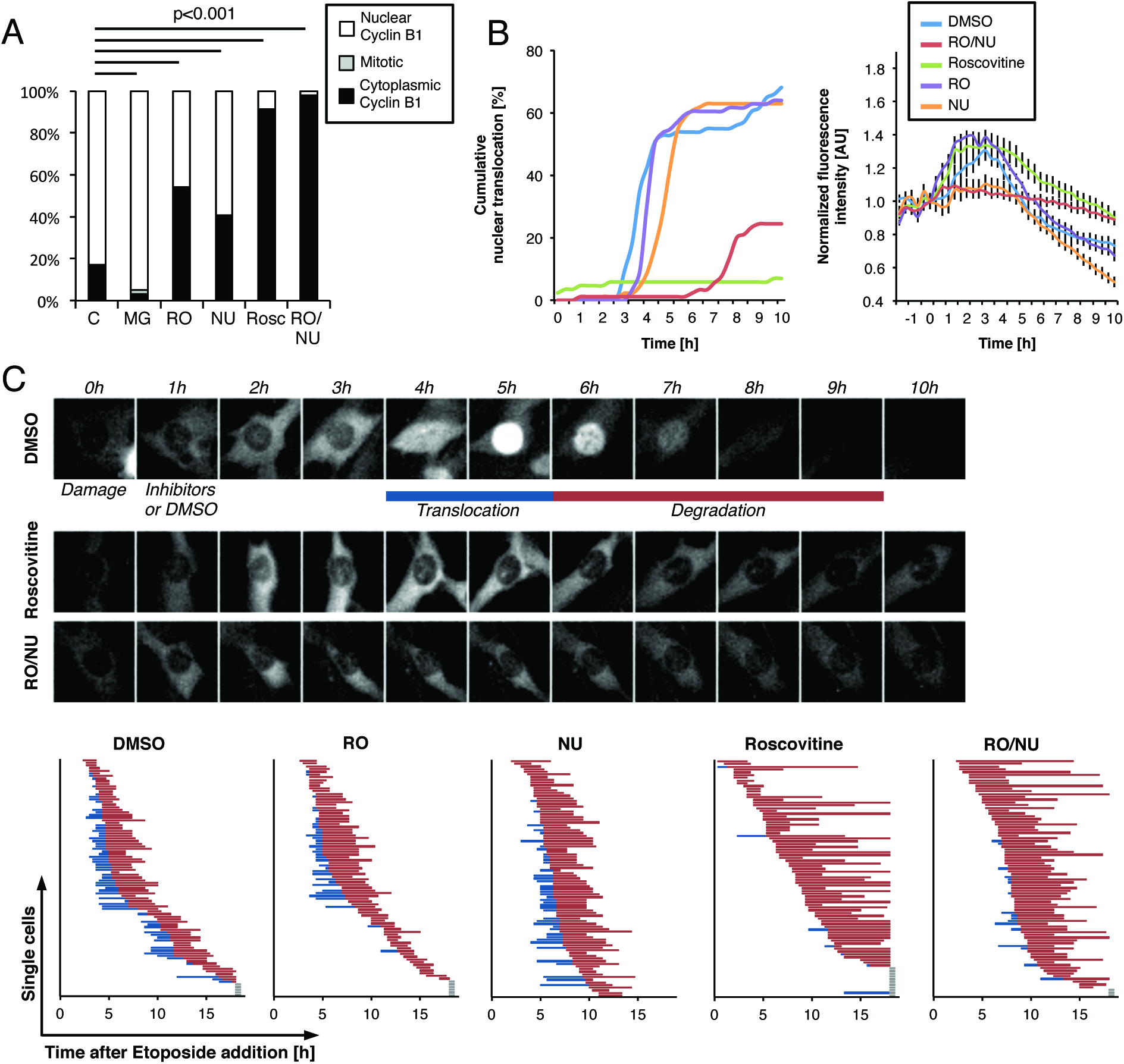
Concerted Cdk1 and Cdk2 activity regulates Cyclin B1 nuclear translocation upon DNA damage. (**A, B, C**) RPE Cyclin B1-eYFP cells were treated with Etoposide at the ‘-1h’ time point. After 1h cells were treated with MG-132, Roscovitine, RO-3306, NU6140, inhibitor combinations as indicated, or mock treated with DMSO (Control). (A) Intracellular localization of Cyclin B1-eYFP was assessed in single cells at 4h after Etoposide addition. Cells were scored to have ‘Increased nuclear Cyclin B1’ if average Cyclin B1 fluorescence in the nucleus was higher than in the cytoplasm. Statistical hypothesis testing was performed using Pearson’s X^2^ test. (B, left graph) Cumulative nuclear translocation was assessed in more than 150 cells for each condition. (B, right graph) Time-lapse microscopy quantifications of Cyclin B1-eYFP levels. The error bars represent standard error of the mean signal of at least eight positions. (C, top panel) Representative images of single cells. (C, graphs) The time point of nuclear translocation and the onset of degradation were determined in single cells. Each line represents a single cell. Blue lines indicate Cyclin B1 nuclear translocation, red lines indicate degradation of Cyclin B1, and grey lines indicate cells with no detectable Cyclin B1 degradation.

DNA-damage induced nuclear translocation of Cyclin B1 in G2 phase marks a decision point for terminal cell cycle exit and senescence [6,11]. Since inhibition of Cdk1/2 activity substantially delayed nuclear translocation of Cyclin B1, we next sought to test if Cdk activity affects whether cells become senescent. To this end, we assessed senescence-associated markers while perturbing Cdk activity using kinase inhibitors or Cdk RNAi. We quantified the occurrence of β-Galactosidase staining (Fig 2A and Supplementary Fig 2A), total and foci-associated staining of H3K9Me2 and HP1b as markers of senescence-associated heterochromatin foci (SAHF) [20], as well as expression of IL-6 as marker for the senescence-associated secretory phenotype (SASP) [21,22] (Fig 2B and Supplementary Fig 2B-D). While long-term treatment with RO-3306 and NU6140 was cytotoxic to cells, we found that all these markers were reduced upon combined Cdk1/2 RNAi or addition of Roscovitine, indicating that Cdk activity stimulates senescence. Similarly, the proliferation capacity, measured by total cell numbers and clonogenic growth, increased after temporal Cdk inhibition (Fig 2C, D and Supplementary Fig 2E). Increased Cdk activity may however not necessarily lead to increased senescence, as we find no evidence that Wee1 inhibition changed senescence-associated markers or proliferation capacity during constant exposure to Etoposide for 4 or 5 days (Fig 2A-D and Supplementary Fig 2B-E). Thus, our data suggest that Cdk activity after DNA damage stimulates induction of senescence, but also that cells contain sufficient Cdk activity to promote senescence during prolonged exposure to a DNA-damaging compound.

**Figure 2.**
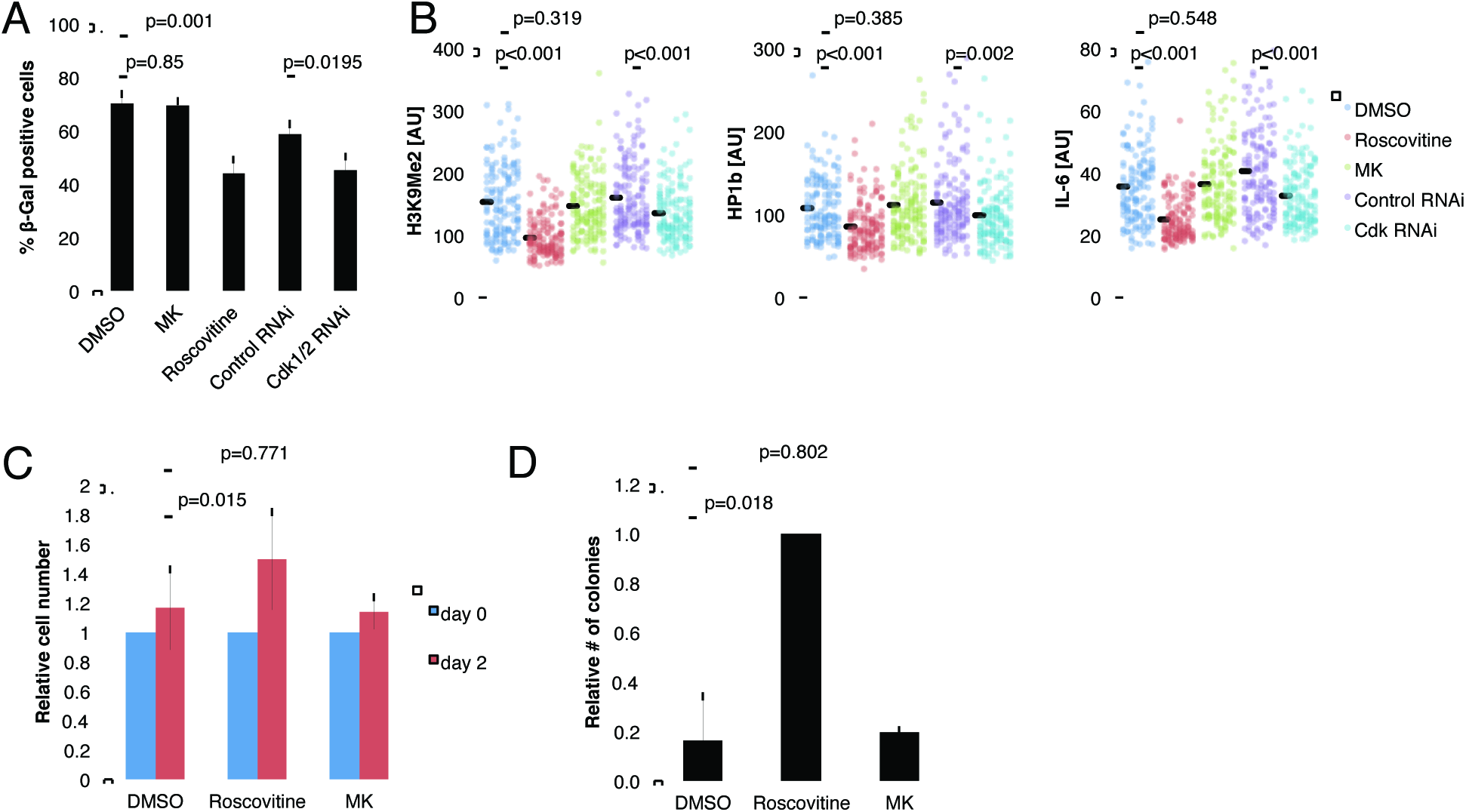
Cdk activity drives senescence induction upon DNA damage. (**A**) Mean and standard deviation from 4 independent experiments of RPE cells treated with Etoposide and after 1h with Roscovitine, MK-1775 (MK) or with DMSO. Alternatively cells were transfected with RNAi for Cdk1 and Cdk2 at 24 and 48h before damage induction in 3 independent experiments. Cells were stained for β-Galactosidase 4 days later. Statistical hypothesis testing was performed using two-sided *t*-test. (B) Quantification of nuclear H3K9Me2, HP1b, and IL-6 levels in RPE cells treated with Etoposide and after 1h with Roscovitine, MK-1775 (MK) or with DMSO. Alternatively cells were transfected with RNAi for Cdk1 and Cdk2 at 24 and 48h before damage induction. Cells were fixed 5 days after damage induction. Statistical hypothesis testing was performed using two-sided *t*-test. (C) Analysis of proliferative capacity. RPE cells were treated with Etoposide and 1h later with Roscovitine, MK-1775 or DMSO. Cells were counted after 5 days, reseeded into fresh medium and counted again after 2 more days. Mean and standard deviation of 3 independent experiments ran in quadruplicates are shown. Statistical hypothesis testing was performed using two-sided *t*-test. (D) Analysis of clonogenic capacity. RPE cells were treated with Etoposide and 1h later with Roscovitine, MK-1775 or DMSO. After 5 days 5000 cells were reseeded into fresh medium and the number of colonies was assessed one week later. Normalized mean and standard deviation of 3 independent experiments ran in quadruplicates are shown. Statistical hypothesis testing was performed using two-sided *t*-test.

### Low levels of Cdk activity are preserved for several hours during a DDR in a cell cycle-dependent manner

Although a DNA damage-mediated checkpoint largely functions by blocking Cdk activity [1,23], our data indicate that Cdk activity is integrated in the DDR. To resolve this apparent paradox, we sought to assess whether Cdk activity persisted after DNA damage. To this end we immunoprecipitated Cyclin A2-eYFP or Cyclin B1-eYFP from gene-targeted RPE cells [6,18] and performed kinase assays on recombinant target proteins that can be phosphorylated by either Cdk2 or Cdk1. Cyclin A2-associated Cdk activity is active through a large part of interphase [24] and was readily detected in a population of unsynchronized cells. Although significantly reduced, Cyclin A2-eYFP associated activity persisted 4h after addition of either Etoposide or the radiomimetic drug Neocarzinostatin (NCS) (Fig 3A). In contrast, Cyclin B1-associated Cdk activity is initially activated at the S/G2 border, and slowly builds up through G2 phase until a dramatic increase initiates mitosis [18,25]. Indeed, we detect a strong Cyclin B1-eYFP associated kinase activity in RPE cells at 10h after release from a thymidine block. However, a markedly reduced Cyclin B1-eYFP associated kinase activity was still present even after 4h Etoposide or NCS treatment, when no mitotic cells were visually detected (Fig 3B). This indicates that although low compared to the activities that initiate mitosis, both Cyclin A and Cyclin B-associated activities are present after DNA damage. Similarly, immunoprecipitated Cdk2 from both unsynchronized and G2 synchronized RPE cells showed reduced but persistent activity 4h after Etoposide treatment (Fig 3C). Thus, Cdk activity can be sustained at a low level after DNA damage in RPE cells.

**Figure 3.**
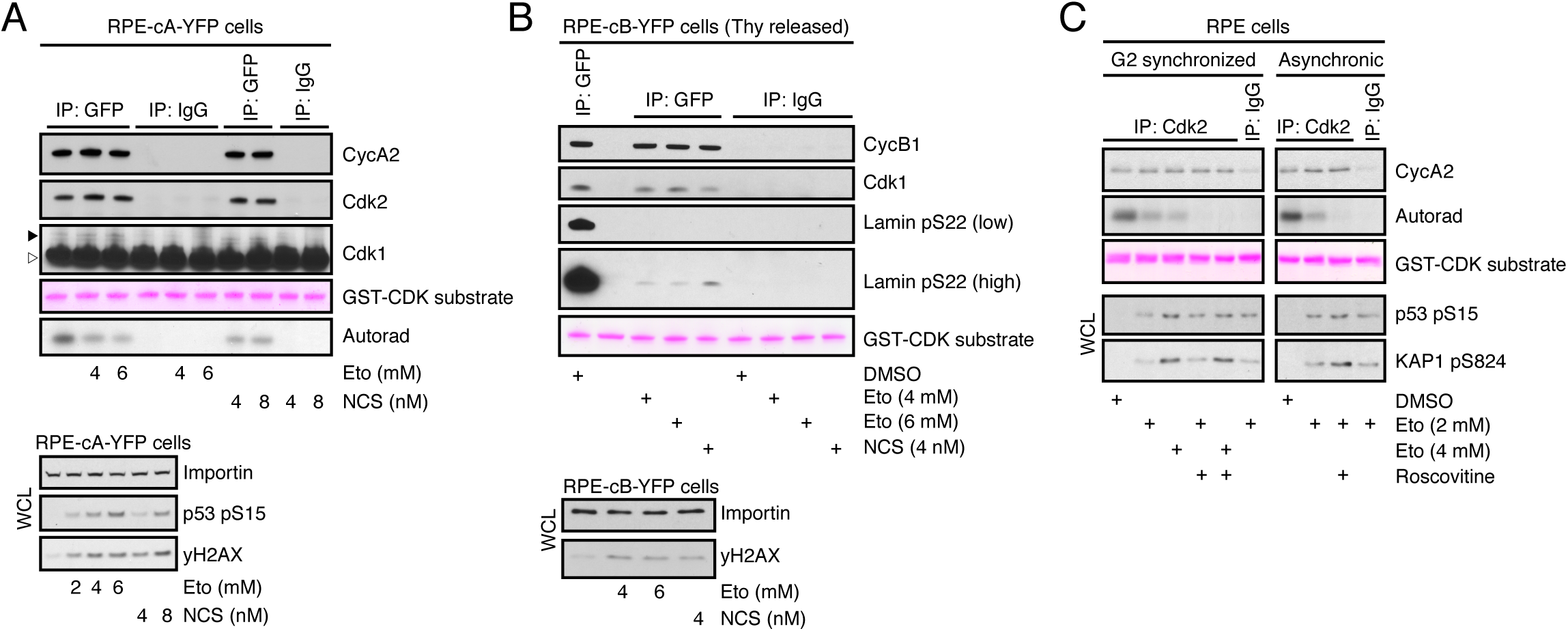
Cyclin A and Cyclin B-associated Cdk activity is preserved during a DNA damage response. (**A**) Asynchronous growing RPE Cyclin A2-eYFP cells were treated with Etoposide or NCS for 4h, lysed and immunoprecipitated with anti-GFP or control antibody. Kinase assay was performed using GST-Cdk substrate peptide and phosphorylation was detected by autoradiography. Co-immunoprecipitation of Cdk2 and Cdk1 was determined by immunoblotting. Arrowhead shows position of Cdk1, empty arrowhead indicates position of IgG. WCL, whole cell lysate. (B) RPE Cyclin B1-eYFP cells were released for 6h from a thymidine block, treated with Etoposide or NCS for 4h, lysed and immunoprecipitated with anti-GFP or control antibody. Kinase assay was performed using GST-LAMS22 substrate peptide and phosphorylation was detected by antibody against Lamin A/C phosphorylated at Ser22. WCL, whole cell lysate. (C) RPE cells were released for 6h from a thymidine block, treated with Etoposide for 4h, lysed and immunoprecipitated with anti-Cdk2 or control antibody. Kinase assay was performed in the absence or presence of Roscovitine and kinase activity was determined as in (a).

We next sought to assess phosphorylation of endogenous Cdk targets in damaged and unperturbed RPE and U2OS cells. To detect ongoing Cdk-mediated phosphorylation we added Cdk inhibitors during the last hour of a 4h Etoposide treatment. For both cell lines, we detect Cdk-dependent phosphorylations after Etoposide treatment in whole cell populations (Fig 4A and Supplementary Fig 4A-D) as well as in single G2 cells (Fig 4B and Supplementary Fig 4E). Cdk target phosphorylation is still detectable at the time of Cyclin B1 nuclear translocation, but not after 24h of DNA damage, suggesting that Cdk activity is preserved until terminal cell cycle exit (Fig 4C and Supplementary Fig 4F). Notably, Cdk target phosphorylation correlated positively with the levels of the DNA damage marker "H2AX, thus excluding the possibility that only mildly damaged cells retain Cdk activity (Supplementary Fig 4G). Furthermore, "H2AX levels were not affected by RO/NU treatment showing that Cdk inhibition does not result in an overall reduction of DNA damage signaling (Supplementary Fig 4G). To assess the cell cycle distribution of Cdk activity during an ongoing DDR we performed quantitative immunofluorescence for Cdk-dependent Lamin A/C phosphorylation and sorted the cells according to their relative position in the cell cycle [6,18,26]. To control for cell cycle-dependent differences in background signals and target site-specificity we added Cdk inhibitors 1h before fixation. In accordance with recent data on Cdk2 [27], we detected initial Cdk1/2 target phosphorylation already during G1, from where it slowly rose throughout S phase before it rapidly increased at the S/G2 border (Fig 4D, ‘Control’). Strikingly, this cell cycle-dependent pattern of Cdk target phosphorylation was still preserved after 4h of continuous Etoposide treatment, albeit at a lower level (Fig 4D, ‘4h Damage’). Inhibition of Cdk1 and Cdk2 decreased Lamin A/C phosphorylation synergistically throughout interphase, indicating a redundancy between both kinases (Fig 4D, ‘4h Damage’; and Supplementary Fig 4H). In summary, we detect Cdk-dependent target phosphorylations in single cells throughout all cell cycle phases during an ongoing DDR.

**Figure 4.**
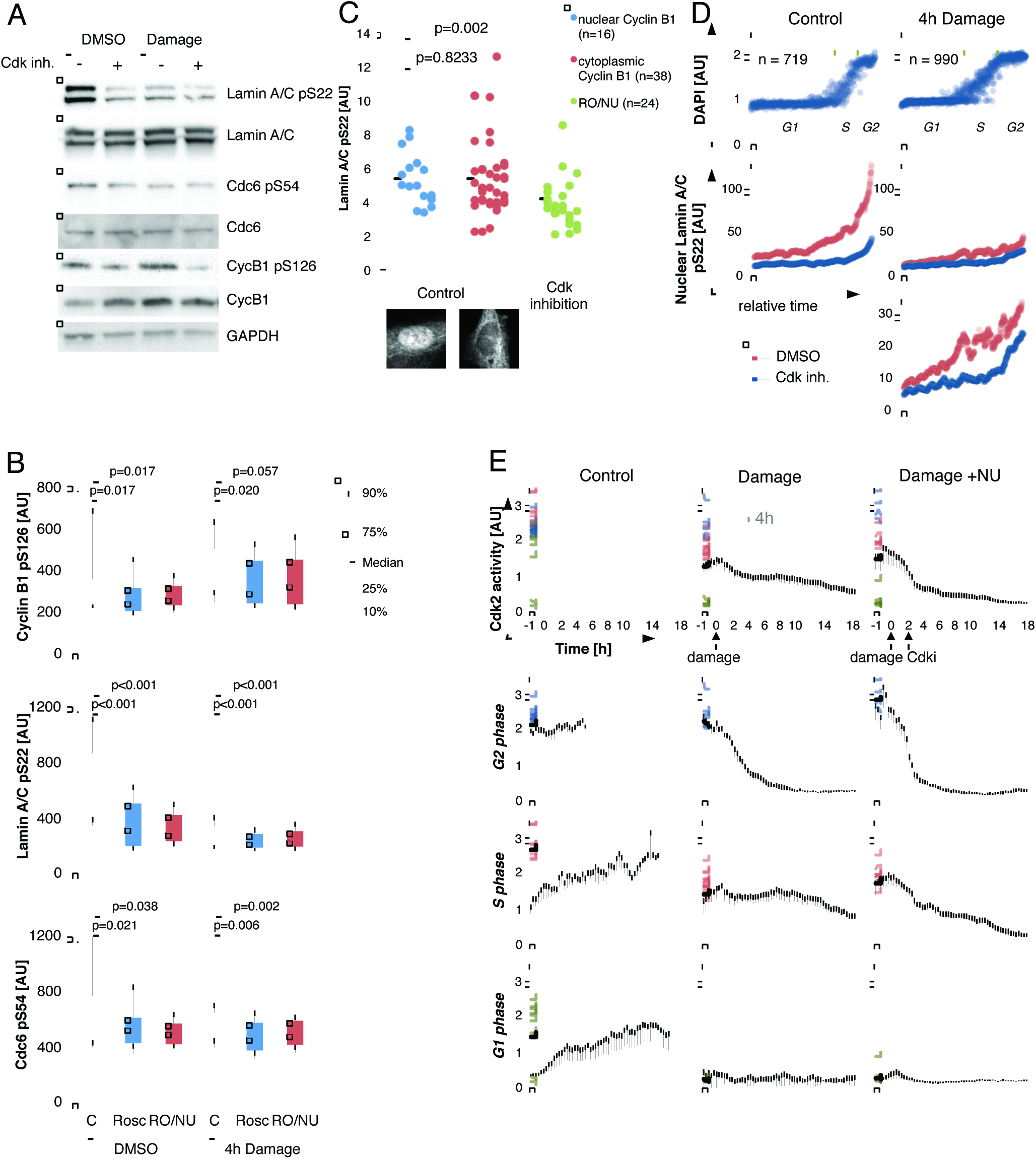
Low levels of Cdk activity are preserved for several hours during a DDR in a cell cycle-dependent manner. (A) Representative Western blot of RPE cells treated with Etoposide or mock treated with DMSO and harvested after 4h. Cells were treated with DMSO (Control) or a combination of Roscovitine, RO-3306 and NU6140 (Cdk inh.) 1h before harvesting. (B) Immunofluorescence quantification of Cyclin B1 pS126, Lamin A/C pS22 and Cdc6 pS54 nuclear fluorescence intensity of interphase cells with 4n DNA content. Cells were treated with Etoposide or mock treated with DMSO and fixed after 4h. Cdk inhibitors were added 1h before fixation. More than 250 cells were analyzed for each condition. G2 cells were identified from DAPI staining. Statistical hypothesis testing was performed using two-sided *t*-test. (C) Quantification of nuclear Lamin A/C pS22 in single G2 RPE cells. Cells were treated as in (B). Cells in G2 phase were identified according to DNA content using DAPI staining. The images show representative single cells with predominantly nuclear or cytoplasmic Cyclin B1. (**D**)Quantification of DNA content (DAPI) and nuclear Lamin A/C pS22 versus estimated time. Cells were sorted for DAPI and Cyclin B1. The Lamin A/C pS22quantifications show a running median of 41 cells. The lower panel shows a zoom-in of the middle panel. Cells were treated with Etoposide or mock treated with DMSO(Control) and fixed after 4h. One hour before fixation, cells were treated with acombination of Roscovitine, RO-3306 and NU6140, or with DMSO. (**E**)Quantification of CDK2 activity in individual, live cells in control and damage conditions. RPE cells expressing a Cdk2 activity probe [27] were followed up to entry into mitosis (Control). The traces were color-coded according to initially low (green),intermediate (red) or high (blue) Cdk2 activity, indicating cells in G1, S or G2 phase respectively [27]. Dashed black lines indicate the respective average. The dashed grey line serves as an indicator of the 4h time point analyzed in fixed-cell experiments.

We next made use of a Cdk2 activity sensor [27] to obtain an independent readout of Cdk2 activity in individual living cells and to gain further insights into the dynamics of Cdk activity during a DDR. In cells with initial high levels of Cdk2 activity, presumably in G2 phase, full inhibition was reached after 6 - 8h of constant Etoposide treatment (Fig 4E, blue tracks). In contrast, cells with initial intermediate levels of Cdk2 activity (suggestive of a S-phase state) sustained this level of Cdk activity for 16h and more (Fig 4E, red tracks). This result is in line with our previous observation that S-phase cells display slower Cyclin B1 accumulation upon DNA damage [6]. Taken together, our data show that Cdk activity is maintained after DNA damage and that the duration and extent of sustained Cdk activity depends on the cell cycle position when DNA damage occurred.

### Cdk activity during DNA damage promotes p21 production

Cell cycle exit and senescence from G2 phase after DNA damage depends on p53 and its transcriptional target p21 [5,6,28,29]. We therefore sought to investigate whether Cdk activity could enhance p53 and p21 expression. In contrast to this hypothesis, we found p53 expression to be elevated when Cdk1 and Cdk2 were inhibited or knocked down, suggesting that Cdk activity does not enhance p53 levels (Fig 5A, B and Supplementary Fig 5A). Cdk inhibition also led to a slight increase in ATM-target phosphorylation, which might contribute to the observed elevated p53 levels (Supplementary Fig 5B, C). Strikingly however, despite an increase in p53, p21 induction was reduced upon Cdk inhibition or depletion, suggesting that the remaining Cdk activity during a DDR promotes p21 expression (Fig 5A, B and Supplementary Fig 5A, D). Further, expression of a constitutively active Cdk1-AF mutant [30] lead to increased p21 levels, supporting a role for Cdk activity to promote p21 expression (Fig 5C). Our data thus suggests that blocking Cdk activity during DNA damage results in an increase in p53 expression, but a decrease in p21 expression. These data are somewhat surprising, given that p21 is a well-established transcriptional target of p53 [31].

**Figure 5.**
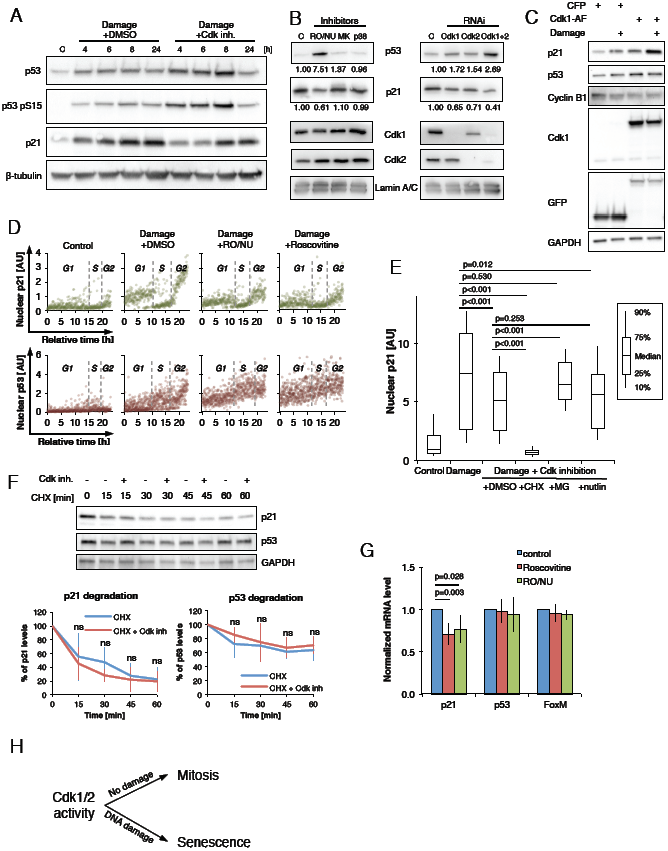
Cdk activity during DNA damage promotes p21 production. (A) Representative Western blot of RPE cells treated with Etoposide and with a combination of Roscovitine, RO-3306 and NU6140 (Cdk inh.) or mock treatment with DMSO 1h later. Cell lysates were prepared at the indicated time points. C, control. (B) Representative Western blots of RPE cells treated with Etoposide and with Roscovitine (Cdk1/2), MK1775 (Wee1), SB202190 (p38) or mock treatment with DMSO 1h later (left blot). Alternatively cells were transfected with the indicated siRNA at 24 and 48h before damage induction (right blot). C, Control. (C) Representative Western blots of RPE cells transfected with Cdk1AF-GFP or control, with and without 4h Etoposide treatment. (D) Quantification of nuclear p21 levels and nuclear p53 level versus estimated time. Cells were sorted for DAPI and Cyclin B1. Cells were treated with Etoposide at 1 µM concentration or mock treated with DMSO (control). Roscovitine, a combination of RO-3306 and NU6140, or DMSO was added 1h after Etoposide treatment. Cells were fixed after 4h. More than 350 cells were analyzed for each condition. (**E**)Immunofluorescence quantification of nuclear p21 levels in G2 cells. RPE cellswere treated with Etoposide and 1h later with Roscovitine (Cdk inhibition) in combination with the indicated drug (DMSO, CHX, MG, nutlin). Cells were fixed 4hafter damage and G2 cells were identified according to DNA content using DAPI staining. (**F**)Representative Western blot of RPE cells treated with Etoposide at time point 0and 1h later with Cycloheximide alone, or Cycloheximide in combination with Cdk inhibition (Roscovitine, RO-3306 and NU6140 - Cdk inh.). Cell lysates were prepared at the indicated time points. Below, quantification of p21 and p53degradation kinetics. The average and standard deviation of three independent experiments are shown. (**G**)RPE cells were treated with Etoposide and 1h later with Roscovitine, acombination of RO-3306 and NU6140, or mock treatment with DMSO. Means and standard deviation of RT-qPCR measurements obtained in 4 independent experiments are shown. Statistical hypothesis testing was performed using two-sided *t*-test. (H) Cdk activity determines cell fate decisions towards mitosis in unperturbed conditions or towards senescence upon DNA damage.

To further investigate Cdk-dependent p21 induction, we next focused on the determinants of p21 expression. Although p21 was induced by Etoposide addition, p21 levels in single cells correlated poorly with the levels of "H2AX staining, indicating that p21 expression is not solely regulated by the amount of damaged DNA present in a cell (Supplementary Fig 5C). In contrast, p21 levels after Etoposide addition showed a strong cell cycle-dependency: p21 was expressed in all cells in G1 phase, virtually absent in cells in S phase [32], and dramatically increased in cells that had crossed the S/G2 border (Fig 5D). Inhibition of Cdk1 and Cdk2 decreased p21 levels both in cells in G1 and G2 phase, indicating that Cdk activity affects p21 expression throughout the cell cycle (Fig 5D, top panel). Analyzing p53 levels, we found stronger induction in S and G2-compared to G1-phase cells. When Cdk activity was inhibited, p53 levels were elevated in all cell cycle phases, again reaching the highest expression in S and G2 phase cells (Fig 5D bottom panel). Thus, our data indicate that the expression levels of p53 and p21 are regulated through the cell cycle and that Cdk activity decreases p53 levels but increases p21 levels in all cell cycle phases.

We next sought to test if Cdk activity affects production or degradation of p21. Addition of the proteasome inhibitor MG-132 and the protein translation inhibitor Cycloheximide affected p21 levels in Etoposide treated G2 cells, suggesting that p21 is continuously produced and degraded. Combined Cdk inhibition and proteasome inhibition resulted in lower p21 levels than in control Etoposide treated cells, suggesting that Cdk-mediated p21 expression cannot be explained solely by differences in p21 degradation (Fig 5E). In line with this finding, Cdk activity did not significantly affect p21 stability after DNA damage (Fig 5F). In contrast, we detect a 20 - 30%-decrease in p21 mRNA levels after Cdk inhibition in Etoposide treated cells (Control) (Fig 5G and Supplementary Fig 5E). Thus, Cdk activity stimulates p21 production at least partly by increasing the amount of p21 mRNA.

## Discussion

A paradox in the cellular response to DNA damage is that a checkpoint is enforced by inhibiting Cdk activity, whereas numerous reports implicate that Cdk activity is needed in a DNA damage response for DNA replication, homologous recombination, and DNA repair [33–38]. Here we show that although the DDR-mediated inhibition of Cdk activity is efficient, low levels of Cdk activity are maintained until terminal cell cycle exit. We suggest that these levels are sufficient to continue Cdk-dependent cell cycle functions such as DNA replication [39] and accumulation of mitotic cyclins [6,40], as well as to promote DNA repair [41], and to maintain the cellular competence for checkpoint recovery [42,43]. Maintaining low levels of Cdk activity during a DDR could provide a necessary time window of slow cell cycle progression in which repair and eventual recovery is possible.

While the remaining Cdk activity sustains important cellular functions during a DDR, it also poses a risk for genome stability. If cells with damaged DNA progress into G2 phase they need to be prevented from entering mitosis, which otherwise would result in chromosome missegregation and propagation of mutations. Indeed, in the absence of p53 or p21, a cell cycle arrest in G2 phase is eventually overrun [8]. We show that Cdk activity is coupled to negative feedback by inducing p21 expression, leading to subsequent Cyclin B1 nuclear translocation and terminal cell cycle exit. As increasing Cdk activity drives mitotic entry, the incorporation of Cdk activity as a positive regulator of p21 expression provides an elegant mechanism to ensure cell cycle exit and senescence when most needed. Thus, our data highlights the overall importance of Cdk activity as cellular read-out for cell cycle position [44,45] and as a regulator of key cell fate decisions (Fig 5H).

The acquisition and maintenance of a G2 DNA damage arrest is very different in transformed and untransformed cells, largely due to misregulation of p53 and p21. This provides both chances and challenges for the selective treatment of cancer cells. Cdk inhibitors in general and Roscovitine (Seliciclib) in particular have shown promising results as part of a combined cancer therapy [46,47]. The antitumor activity and specificity of Cdk inhibitors is mainly attributed to the increased requirement of Cdk activity in highly proliferating tumor cells. In addition, Cdk inhibitors have been proposed as drug candidates in combination with radiation therapy or chemotherapy [48]. Our results raise some caution as they suggest that inhibition of Cdk activity during DNA damage interferes with checkpoint signaling, namely p21 expression and induction of senescence. This could lead to increased checkpoint override in the presence of DNA damage and potentially to sustained proliferation of damaged non-tumor cells.

We propose that upon DNA damage, a low level of cell cycle-dependent Cdk activity is retained. This activity exerts an important dual role within the DNA damage checkpoint response. In S phase Cdk activity allows cells to continue cell cycle progression, DNA replication, and repair. In G2 phase Cdk activity promotes p21 production, creating a strong negative feedback loop that constitute a logical gate to ensure a robust decision towards terminal cell cycle exit. Interestingly, mutations in oncogenes frequently result in increased Cdk activity, thereby driving cell proliferation, and inducing DNA replication stress [49]. Our data suggests that increased Cdk activity in combination with replication stress would lead to increased cell cycle exit from G2, which is in line with the robust senescence block that is observed as an early barrier for tumor progression [50].

## Methods

### Cell culture

RPE and U2OS cell lines were cultured in an ambient-controlled incubator at 37 °C with 5% CO_2_ in Dulbecco’s modified eagle medium (DMEM)/Nutrient mixture F-12 (DMEM/F-12) + GlutaMAX (Invitrogen) supplemented with 10% FBS (FBS, HyClone) and 1% Penicillin/Streptomycin (Pen/Strep; HyClone), and DMEM + GlutaMAX (Invitrogen) supplemented with 6% heat-inactivated fetal bovine serum and 1% Pen/Strep respectively. For live-cell imaging experiments cells were cultured in Leibowitz’s L-15 medium (Invitrogen) supplemented with 10% FBS and 1% Pen/Strep.

### Plasmids, cloning, purification and transfection

The use of the live-cell sensor for Cdk2 activity (kindly provided by Tobias Meyer and Sabrina Spencer) has been described previously [27]. For experiments involving constitutively active Cdk1, cells were transfected using 2.5 µg Cdk1AF-GFP (kindly provided by Rob Wolthuis) or ECFP1-C1 (Clontech) using Lipofectamine 2000 (Invitrogen) 24h before analysis of the phenotype.

To clone substrates for the kinase assays, DNA fragments corresponding to the optimal Cdk2 substrate peptide HHASPRK or STPLSPTRIT peptide derived from Lamin A were ligated in frame into pGEX6P plasmid [51]. GST, GST-CDK2 or GST-LAMS22 substrates were purified from BL21 bacteria induced by 0.5 mM IPTG for 5h using glutathione beads.

### Inhibitors, and RNAi

The inhibitors used in this study were employed at the following concentrations: Roscovitine at 25 µM (Cdk inhibitor; Selleck Chemicals), NU6140 at 10 µM (Cdk2 inhibitor; Calbiochem), RO-3306 at 10 µM (Cdk1 inhibitor; Calbiochem), MG-132 at 10 µM (Inhibitor of the proteasome; Sigma Aldrich), BI2536 at 100 nM (Plk1 inhibitor; Selleck Chemicals), MK-1775 at 100 µM (Wee1 inhibitor; Selleck Chemicals), Nutlin-3 at 13 µM (Mdm2 antagonist; Sigma Aldrich), SB202190 at 10 µM (p38 inhibitor; Selleck Chemicals), Cycloheximide at 10 µg/ml (Inhibitor of protein translation; Sigma Aldrich). Etoposide (topoisomerase II inhibitor; Sigma Aldrich) was employed at 1 µM, which is sufficient to induce robust checkpoint arrest in cells at all cell cycle stages [6]. Supplementary figure 1d shows the robustness of the checkpoint arrest as well as the efficiency of Cdk inhibitor treatment. SMARTpool ON-TARGET plus siRNAs targeting CDKN1A (p21), CDK1, or CDK2 were purchased from Dharmacon and employed at a concentration of 20 nM using HiPerFect (Qiagen) transfection at 48h and 24h before analysis of the phenotype.

### Live-cell microscopy and quantitative immunofluorescence

Live-cell imaging experiments were done as previously described [6]. For quantitative immunofluorescence cells were fixed and immunostained as previously described [6]. Images were acquired on an ImageXpress system (Molecular Devices) using a 40x NA 0.6 or a 60x NA 0.7 objective. Images were manually screened for aberrant staining or illumination, and processed and analyzed using CellProfiler and ImageJ. Background subtraction and image analysis for the identification of cell nuclei was essentially done as previously described [52]. Supplementary figure 4i shows imaging examples and analysis including several controls. To assess kinetics from quantitative immunofluorescence cells were ordered based on increasing, median-normalized DAPI and/or nuclear Cyclin B1 fluorescence [18,26]. The cells were linearly distributed according to their fluorescence value between the time 0h and 23h. The approximate borders between cell cycle phases were visually identified according to the DAPI profile [18,26]. Foci analysis was performed by subtracting a Gaussian filter blurred image from the original image, and measuring the integrated intensity of the resulting foci per nucleus using CellProfiler. The following antibodies were used: Cyclin B1 V152 (1:400; #4135 Cell Signaling), γH2AX (1:400; #9781 Cell Signaling), p53 7F5 (1:200; #2527 Cell Signaling), p21 12D1 (1:400; #2947 Cell Signaling), Lamin A/C pS22 (1:400; #2026 Cell Signaling), Cyclin B1 pS126 (1:200; ab55184 abcam), Cdc6 pS54 EPR759Y (1:200; ab75809 abcam), IL6 (1:1000; ab9324 abcam), H3K9Me2 (1:500; ab1220 abcam), HP1b 1MOD-1A9 (1:1000; MAB3448 Millipore).

### Immunoblotting

The following antibodies were used: Lamin A/C pS22 (1:400; #2026 Cell Signaling), Cdc6 pS54 EPR759Y (1:200; ab75809 abcam), Cyclin B1 pS126 (1:200; ab55184 abcam, 1:100; ab3488 abcam), p53 DO-1 (1:500; sc-126 Santa Cruz), p53 pSer15 (1:200, #9284 Cell Signaling), p21 12D1 (1:1000; #2947 Cell Signaling), β-tubulin 9F3 (1:1000; #2128S Cell Signaling), Cdk1 POH1 (1:1000; #9116 Cell Signaling), Cdk1 (1:200; HPA003387 Atlas antibodies), Cdk2 78B2 (1:1000; #2564 Cell Signaling), Lamin A/C (1:2000; #4777 Cell Signaling), GAPDH (1:15000-25000; G9545 Sigma Aldrich), pKap1 (1:500; A300-767A Bethyl antibodies) and pChk2 (1:1000; #2661 Cell Signaling).

### qPCR

For RT-qPCR total RNA was extracted from the cells with an RNeasy Mini kit (Qiagen). Reverse transcriptase reaction was set up using 250 ng of total RNA and 10 pmol oligo (dT) primers. The qPCR reactions were set up using Fast SYBR Green Master Mix (life technologies). Samples were run on a 7500 Fast Real-Time PCR System (life technologies) using the following primers: p21-forward (5'-AGG CAC CGA GGC ACT CAG AG-3'), p21-reverse (5-AGT GGT AGA AAT CTG TCA TGC TG-3), p53-forward (5'-ATG GAG GAG CCG CAG TCA GAT-3'), p53-reverse (5'-GCA GCG CCT CAC AAC CTC CGT C-3)', FOXM1-forward (5’-ACC CAA ACC AGC TAT GAT GC-3’), FOXM1-reverse (5’-GAA GCC ACT GGA TGT TGG AT-3’), GAPDH-forward (5'-CGG AGT CAA CGG ATT TGG TCG TAT-3'), GAPDH-reverse (5'-AGC CTT CTC CAT GGT GGT GAA GAC-3'). Primers were obtained from Sigma Aldrich.

### Senescence-associated β-Galactosidase assay, cell proliferation, and clonogenic assay

β-Gal stainings were performed using the ‘Senescence β-Galactosidase Staining Kit’ (Cell Signaling).

To determine cell proliferation potential, cells were seeded in quadruplicates in 6-well plates 1 day before treatment with Etoposide with and without different inhibitors. Cells were counted after 4 days of treatment, reseeded into fresh medium, and counted again after an additional 2 days.

To determine clonogenic capacity, cells were treated with Etoposide with and without different inhibitors. After 4 days 5000 cells were seeded into fresh medium in quadruplicates in 6-well plates. After an additional 7 days the samples were fixed with 10% formaline and stained with 0.5% (w/v) crystal violet before the number of colonies was assessed.

### Kinase assay

Asynchronously growing RPE Cyclin A2-eYFP cells or RPE cells released for 6h from thymidine block (2.5 mM, 24h) were treated with DMSO or with indicated doses of Etoposide or Neocarzinostatin for 4h and lysed in ice cold IP buffer (50 mM Hepes pH 7.4, 150 mM NaCl, 10 % glycerol, 0.1 % NP-40) supplemented with protease and phosphatase inhibitors (Roche). Normalized cell extracts were incubated with 2 µg of control IgG, antibody against GFP or Cdk2 for 1h and for additional 1h with protein A/G beads (Pierce). Beads were washed twice with IP buffer and once with kinase buffer (25 mM MOPS pH 7.2, 12.5 mM glycerol 2-phosphate, 25 mM MgCl2, 5 mM EGTA, 2 mM EDTA and 0.25 mM DTT). Beads were incubated with kinase buffer supplemented with 100 μM ATP, 5 μCi ^32^P-γ-ATP and purified GST-Cdk substrate (2 μg) for 20 min at 30°C. Where indicated 12 mM Roscovitine was added to the kinase buffer. Reaction was stopped by addition of 4x Laemli buffer and boiling for 5 min. Proteins were separated by SDS-PAGE and phosphorylation of GST-Cdk substrate was detected by autoradiography. Alternatively, RPE Cyclin B1-eYFP cells were synchronized by thymidine for 24h, released for 6h when cells reach G2 phase and then treated with etoposide for 4h. Immunoprecipitation and kinase assay was performed as mentioned above, except GST-LAMS22 substrate was used. Phosphorylation of GST-LAMS22 was detected by immunoblotting using an antibody against Lamin A/C phosphorylated at Ser22 (see above).

## Statistical analysis

Statistical hypothesis testing for differences between the means of two populations was done using Welch’s *t*-test, an adaptation of Student’s *t*-test, that is reliable for populations with unequal variances and sample sizes, and remains robust for skewed distributions [53].

Statistical hypothesis testing for observed differences in sets of categorical data was done using Pearson’s X^2^ test.

## Acknowledgements

We thank lab members for critical comments on the manuscript. This work was supported by grants from the Swedish Research Council, the Swedish Foundation for Strategic Research, and the Swedish Cancer Society. LM was supported by the Grant Agency of the Czech Republic (13-18392S) and by RVO: 68378050.

## Author contributions

E.M. and A.L. designed the study and wrote the manuscript. E.M., H.SC., and L.M. carried out the experiments. E.M., H.SC., L.M., and A.L. analyzed the data.

---

**Supplementary Figure 1.**
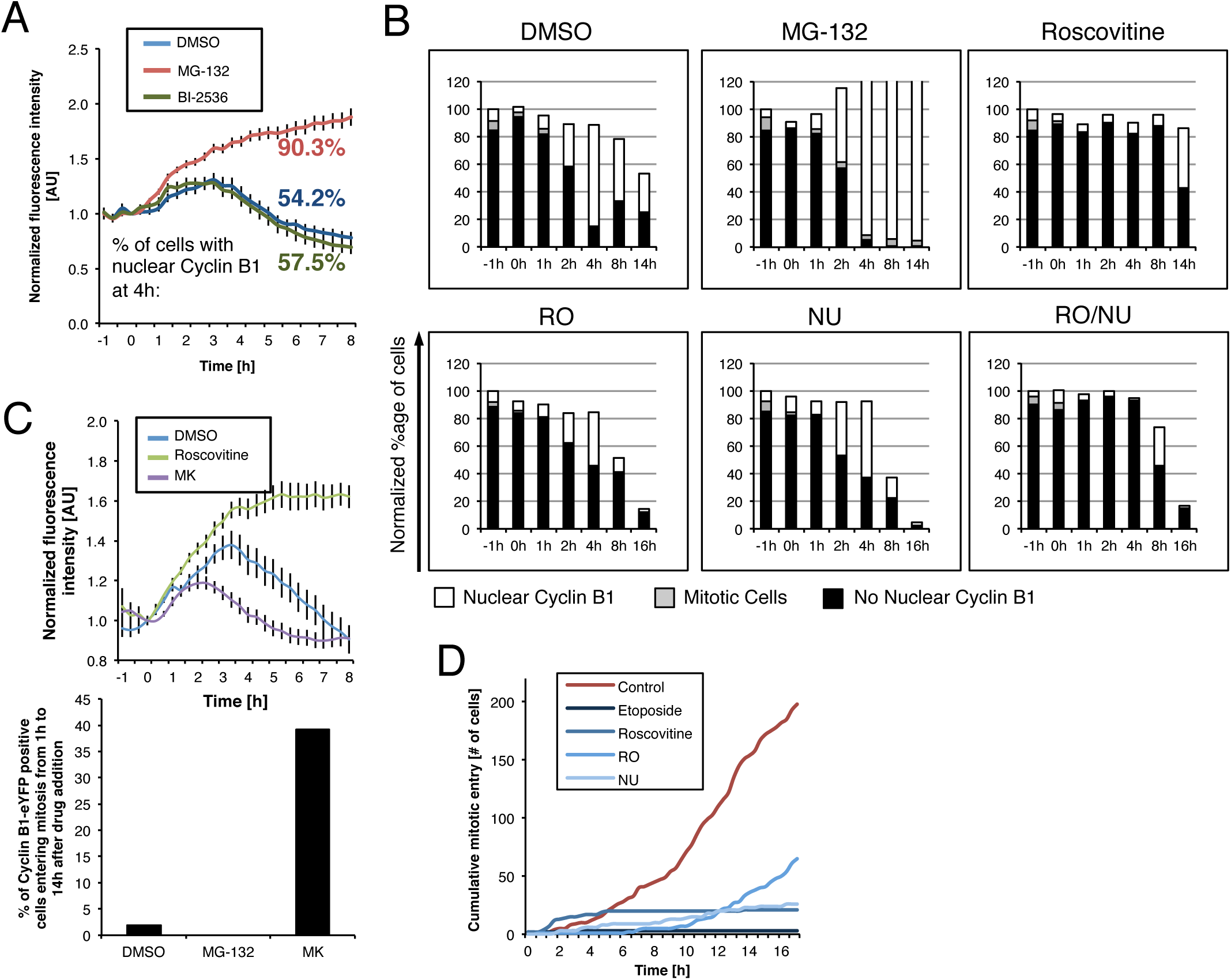
Cdk1 and Cdk2 activity, but not Plk1 regulates Cyclin B1 nuclear translocation upon DNA damage. (A) Plk1 activity does not affect Cyclin B1 degradation. Time-lapse microscopy quantifications of RPE Cyclin B1-eYFP cells treated with Etoposide at time point ‘-1h’. MG-132 or BI-2536 was added at time point ‘0h’. The error bars represent standard error of the mean signal of at least eight positions. The localization of Cyclin B1-eYFP was assessed in more than 120 cells for each condition and the percentage of cells with predominantly nuclear Cyclin B1 at the ‘4h’ time point is indicated. (B) RPE Cyclin B1-eYFP cells were treated with Etoposide at the ‘-1h’ time point. After 1h cells were treated with MG-132, Roscovitine, MK-1775, RO-3306, NU6140, or inhibitor combinations as indicated. The intracellular localization of Cyclin B1-eYFP was assessed at different time points starting with more than 180 single cells for each condition at time point ‘-1h’. The values were normalized to the initial cell number. (C) Increased Cdk activation causes earlier degradation of Cyclin B1 and increased slippage of the DNA damage checkpoint. Time-lapse microscopy quantifications of Cyclin B1-eYFP levels (upper graph) and mitotic progression (lower graph) in RPE Cyclin B1-eYFP cells. Etoposide was added at the ‘-1h’ time point. After 1h cells were treated with Roscovitine, or MK-1775. The error bars represent standard error of the mean signal of at least eight positions (upper graph). Mitotic progression was analyzed in more than 200 cells for each condition (lower graph). (D) Cumulative mitotic entry was scored in RPE Cyclin B1-eYFP cells treated with DMSO (‘control’), Etoposide, Roscovitine, RO-3306, or NU6140 to test for the efficiency of inhibition.

**Supplementary figure 2.**
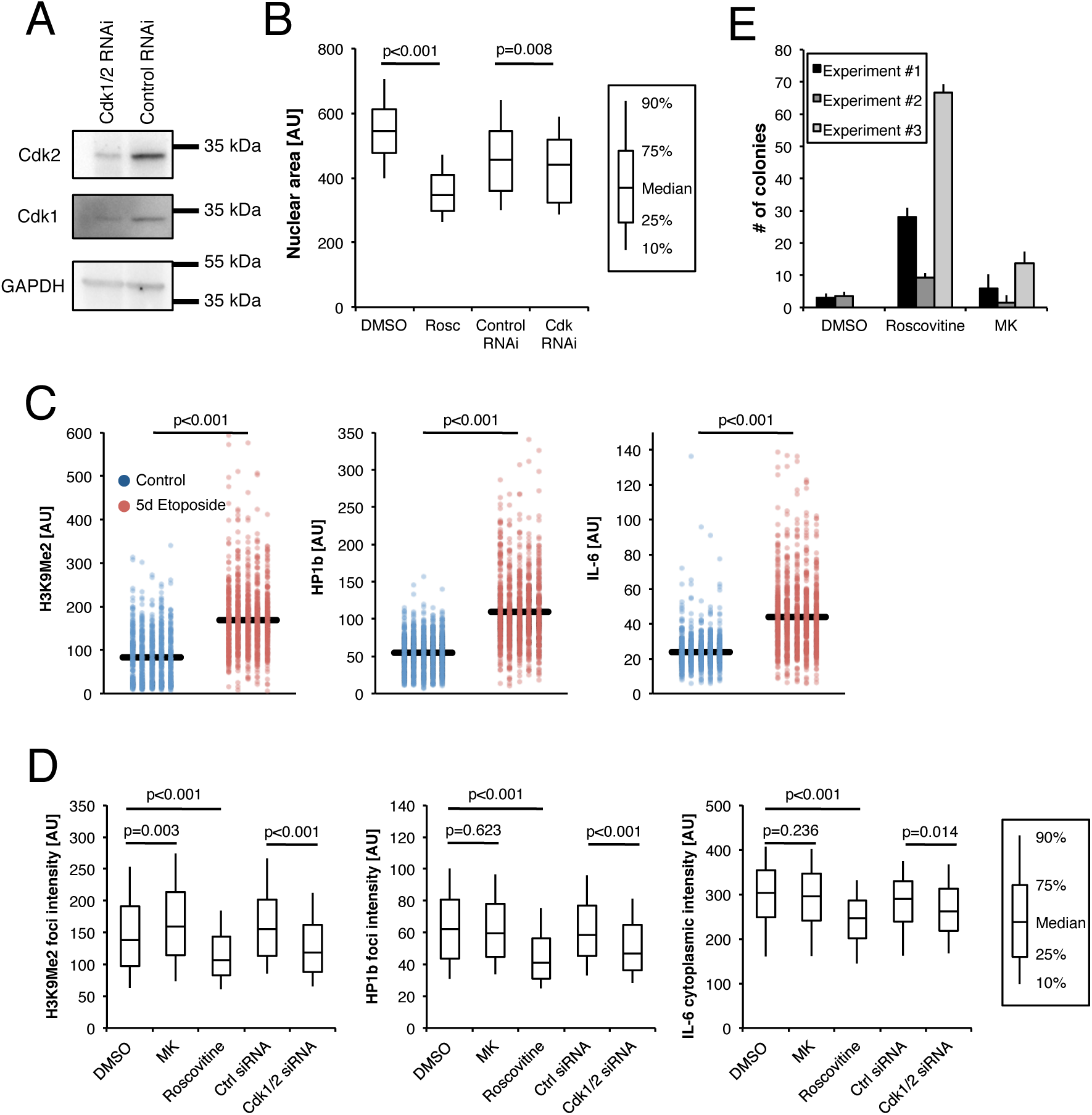
Cdk activity induces senescence upon DNA damage. (A) Western blot of a typical siRNA knockdown of Cdk1 and Cdk2 in a 96-well format. Cells were transfected with RNAi for Cdk1 and Cdk2 24 and 48h before sample preparation. (B) Increased nuclear size as an indicator of cellular senescence [54]. RPE cells were treated with Etoposide. Roscovitine or DMSO were added 1h later. Alternatively, cells were transfected with siRNA for Cdk1 and Cdk2 at 24 and 48h before damage induction. Cells were fixed 4 days after damage induction, stained with DAPI and nuclear size was assessed in more than 250 cells for each condition. Statistical hypothesis testing was performed using two-sided *t*-test. (C) Quantification of nuclear H3K9Me2, HP1b, and IL-6 levels in control RPE cells and cells treated with Etoposide for 5 days. Statistical hypothesis testing was performed using two-sided *t*-test. (D) Quantification of nuclear foci intensity of H3K9Me2 and HP1b, and cytoplasmic IL-6 levels. RPE cells treated with Etoposide and after 1h with Roscovitine, MK-1775 (MK) or with DMSO. Alternatively cells were transfected with siRNA for Cdk1 and Cdk2 at 24 and 48h before damage induction in 3 independent experiments. Cells were fixed 5 days after damage induction. Statistical hypothesis testing was performed using two-sided *t*-test. (**E**)Colony formation capacity of RPE cells treated with Etoposide and DMSO,Roscovitine (Rosc) or MK-1775. Data from 3 independent experiments are shown.

**Supplementary figure 4.**
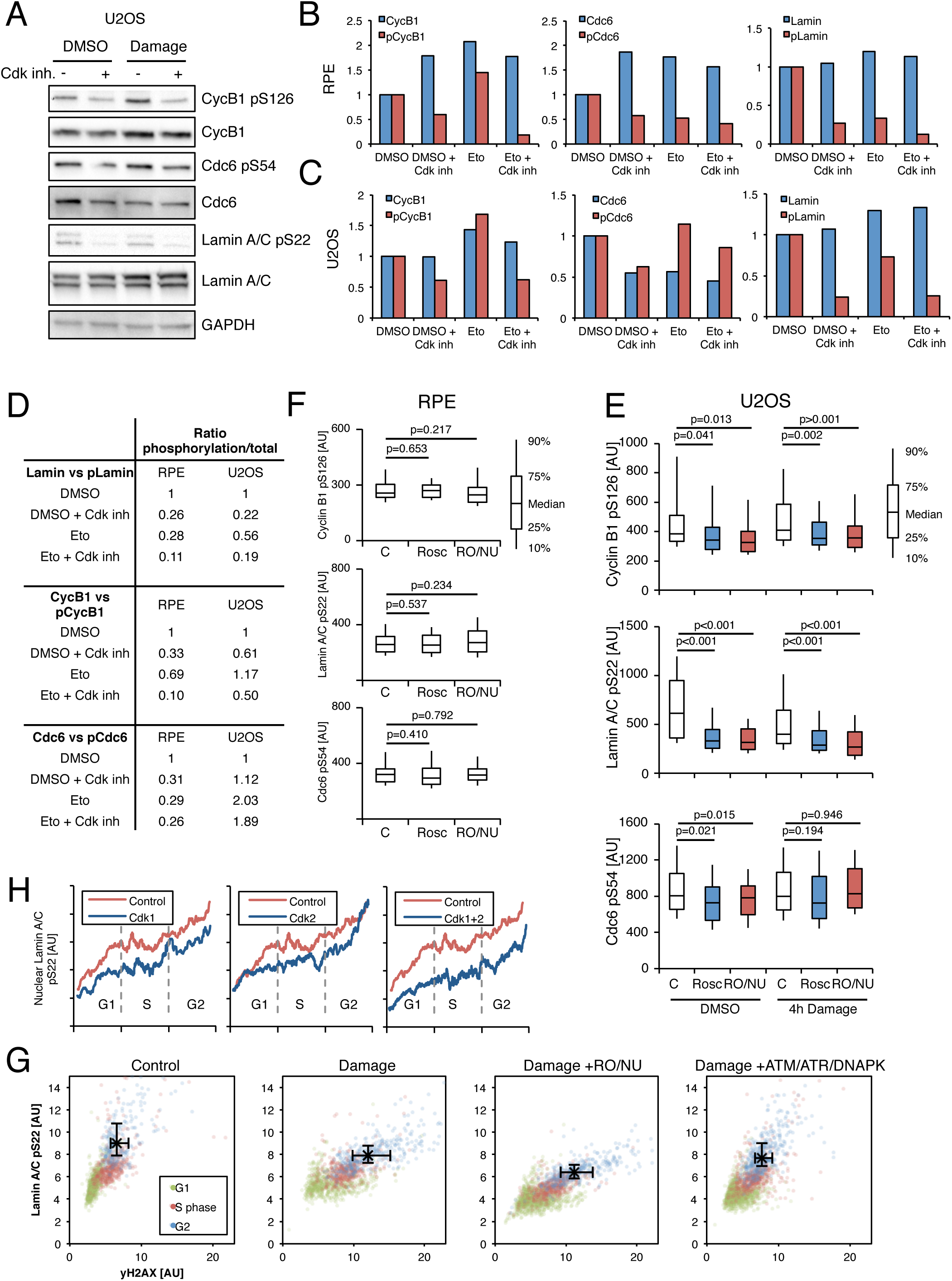
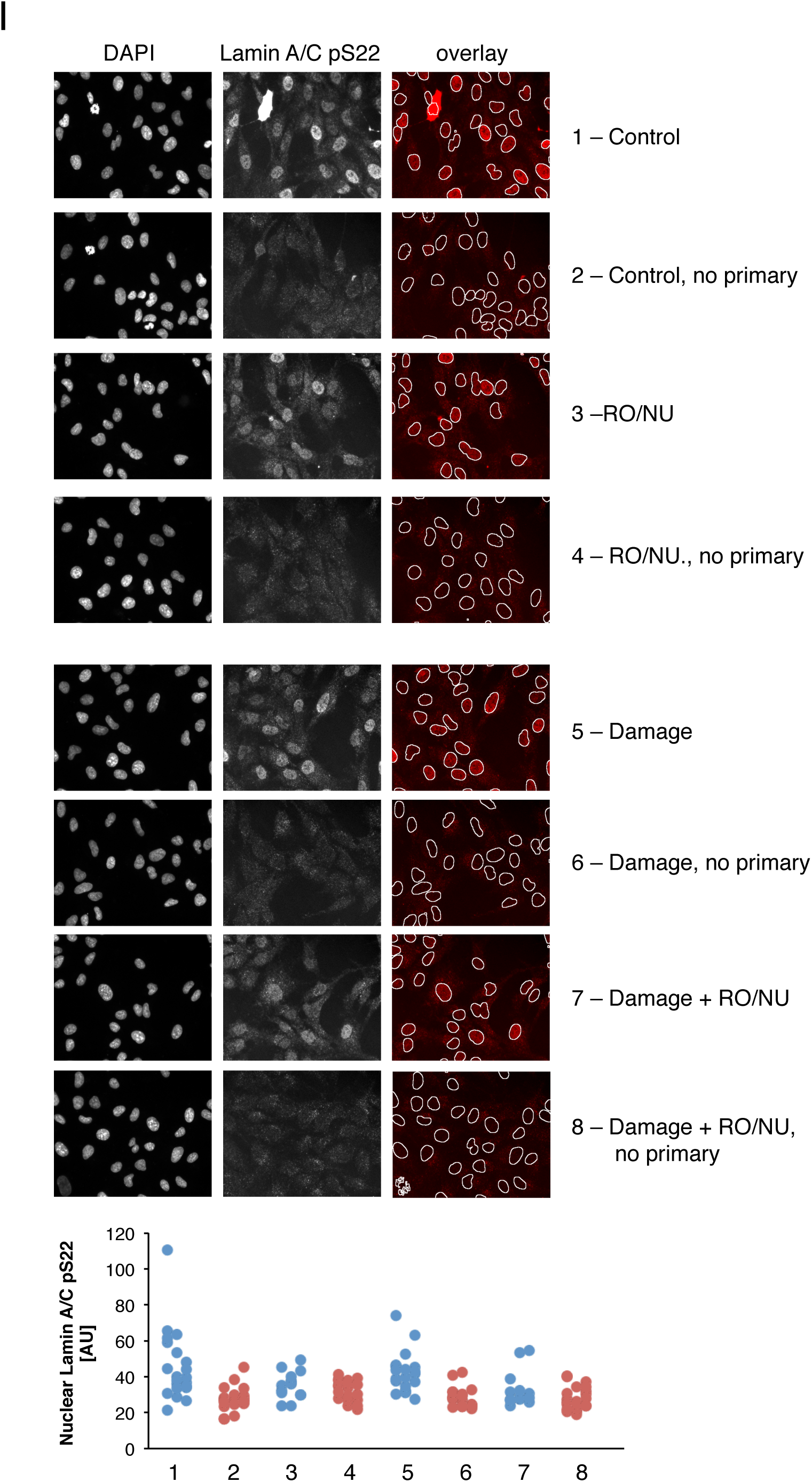
dk activity is retained after DNA damage in RPE and U2OS cells. (A) Representative Western blot of U2OS cells treated with Etoposide or mock treated with DMSO and harvested after 4h. One hour before harvest cells were treated with a combination of Roscovitine, RO-3306 and NU6140 (Cdk inh.). (B) Quantification of Western blots shown in (Figure 4A). The values were normalized to the DMSO condition for every antibody. (C) Quantification of Western blots in (A). The values were normalized to the DMSO condition for every antibody. (D) Results of the ratio between phosphorylated protein and total protein of the targets quantified in (b) and (c). (**E**)Immunofluorescence quantification of Cyclin B1 pS126, Lamin A/C pS22 and Cdc6 pS54 nuclear fluorescence intensity of interphase 4n DNA content U2OS cells treated with Etoposide or mock treated with DMSO and fixed after 4h. Cdk inhibitors were added 1h before fixation. G2 cells were identified from DAPI staining. Morethan 150 cells were analyzed for each condition. Statistical hypothesis testing was performed using two-sided *t*-test. (**F**)Immunofluorescence quantification of Cyclin B1 pS126, Lamin A/C pS22 and Cdc6 pS54 nuclear fluorescence intensity in interphase RPE cells with 4n DNA content cells treated as in (E) but fixed at 24h. More than 150 cells were analyzed per condition. Statistical hypothesis testing was performed using two-sided *t*-test. (**G**)Immunofluorescence quantification of nuclear Lamin A/C pS22 levels versusnuclear "H2AX levels in RPE cells. Cell cycle phases were identified according toDNA content using DAPI staining. Black bars represent the 25^th^ and 75^th^ percentile ofthe G2 population. The intersection between the bars indicates the median intensity ofthe two stainings in G2 cells. (**H**) Quantification of nuclear Lamin A/C pS22 versus estimated time. RPE cells were sorted for DAPI and Cyclin B1. A running median of 30 cells is shown. Cells were treated with Etoposide and fixed after 4h. One hour before fixation cells were treated with RO-3306, NU6140, a combination of both, or mock treated with DMSO. Cell cycle phases were identified according to DNA content using DAPI staining. More than 500 cells were analyzed for each condition. (**I**) Representative images of cells treated with and without Etoposide (control, damage) and with and without Cdk inhibition (RO-3306 and NU6140) stained with and without primary antibodies. The graph shows the quantification of nuclear Lamin A/C pS22 in the depicted images.

**Supplementary figure 5.**
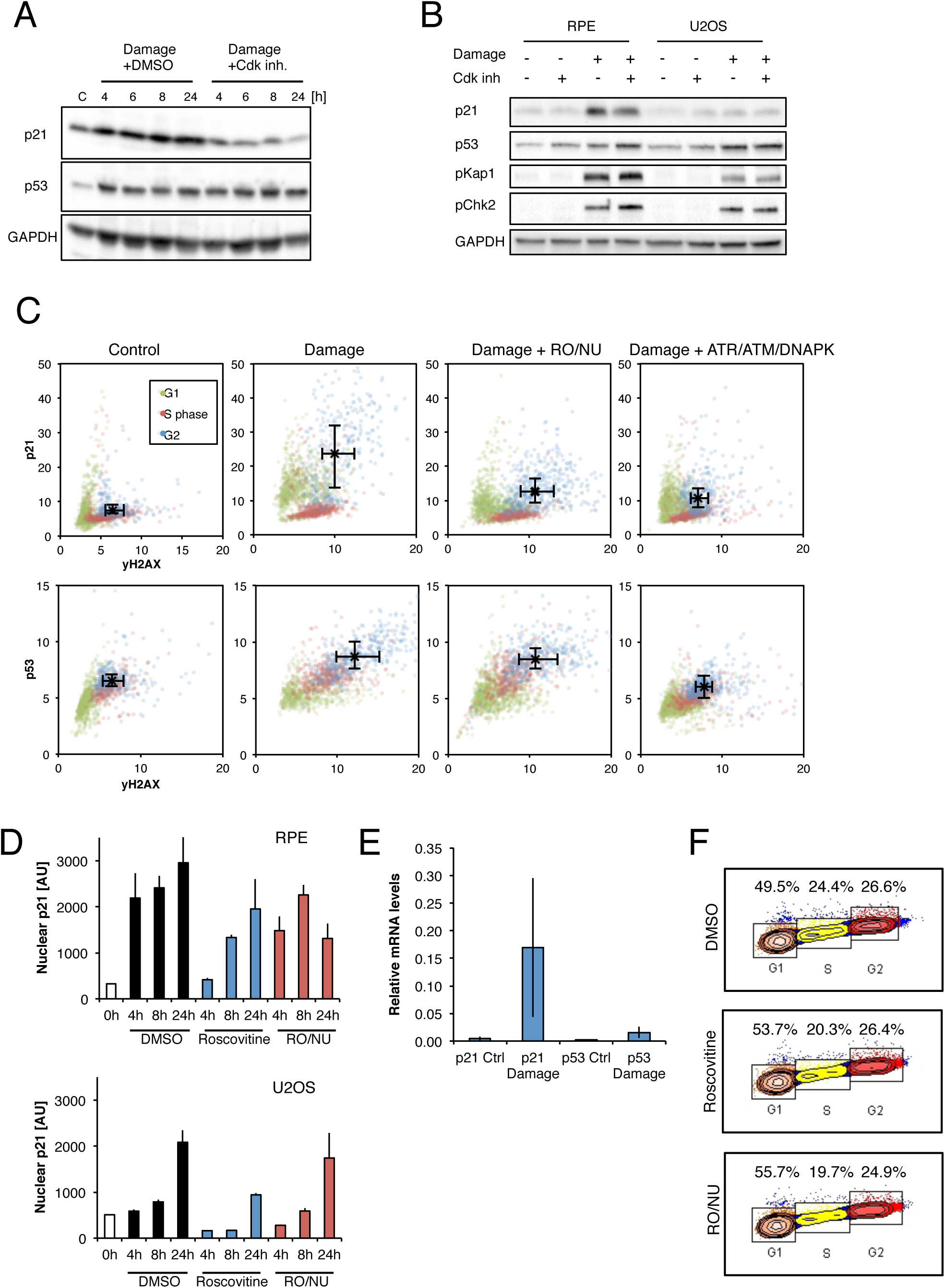
Cdk activity promotes p21 production in RPE and U2OS cells. (A) Representative Western blot of U2OS cells treated with Etoposide (Damage) and 1h later with a combination of Roscovitine, RO-3306 and NU6140 (Cdk inh.) or DMSO. Cell lysates were prepared at the indicated time points. C, Control. (B) Representative Western blot of RPE and U2OS cells treated with or without Etoposide, and with and without Cdk inhibitor (a combination of Roscovitine, RO-3306 and NU6140) 1h later. (C) Immunofluorescence quantification of nuclear p21 and nuclear p53 levels versus nuclear "H2AX levels. Cells were treated with DMSO (Control), or with Etoposide and 1h later with a combination of RO-3306 and NU6140, a combination of ATR, ATM and DNA-PK inhibitor, or DMSO. Cell cycle phases were identified according to DNA content using DAPI staining. Black bars represent the 25^th^ and 75^th^ percentile of the G2 population. The intersection between the bars indicates the median intensity of the two stainings in G2 cells. (**D**)Immunofluorescence quantification of p21 nuclear fluorescence intensity in interphase RPE and U2OS cells with 4n DNA content. Cells were treated with Etoposide and 1h later with Roscovitine, a combination of RO-3306 and NU6140, orDMSO. Cells were fixed at the indicated time points. Mean and standard deviation are shown. More than 100 cells were analyzed for each condition. (**E**)Relative p21 and p53 mRNA levels in Etoposide and untreated RPE cells measured by RT-qPCR. Averages and standard deviations obtained from 3independent experiments are shown. (**F**)Flow cytometry analysis of RPE cells treated with Etoposide and 1h later with Roscovitine, a combination of RO-3306 and NU6140, or DMSO. The cells were fixed after 4h. DNA content was assessed by PI staining.

